# isiKnock: *in silico* knockouts in biochemical pathways

**DOI:** 10.1101/313858

**Authors:** Jennifer Scheidel, Heiko Giese, Börje Schweizer, Leonie Amstein, Jörg Ackermann, Ina Koch

**Affiliations:** Molecular Bioinformatics, Institute of Computer Science, Johann Wolfgang Goethe-University Frankfurt am Main, Robert-Mayer-Straße 11-15, 60325 Frankfurt am Main, Germany

**Author notes:** Shared first author (Jennifer Scheidel, Heiko Giese).

## Abstract

**Summary:** isiKnock is a new software that automatically conducts *in silico* knockouts for mathematical models of biochemical pathways. The software allows for the prediction of the behavior of biological systems after single or multiple knockout. The implemented algorithm applies transition invariants and the novel concept of Manatee invariants. A knockout matrix visualizes the results. The tool enables the analysis of dependencies, for example, in signal flows from the receptor activation to the cell response at steady state.

**Availability and Implementation:** isiKnock is an open-source tool, freely available at http://www.bioinformatik.uni-frankfurt.de/tools/isiKnock/index.php. It requires at least Java 8 and runs under Microsoft Windows, Linux, and Mac OS.

## 1 Introduction

Biochemical pathways are highly intertwined processes. The perturbation of pathway components can reveal their complicated interplay and dependencies. The computational analysis of the knockout behavior is a valuable approach to predict the system’s behavior for knockout conditions and to detect inconsistencies in the mathematical model caused, e.g., by incomplete data. The theoretical analysis can provide new insights to the system’s behavior and easily simulate hypotheses.

In Scheidel *et al.,* 2016, we have introduced the concept of *in silico* knockout analysis at steady state. The analysis is based on transition invariants, also known as elementary modes (Schuster *et al.,* 2000), describing the complete set of basic processes of the model (Koch *et al.*, 2011). We have demonstrated the benefit of the new concept for a key defense mechanism against the pathogen *Salmonella Typhimurium* called xenophagy. We have compared the simulated knockouts with available data from experimental perturbation studies to ensure the credibility of the theoretical network of the xenophagic capturing of *Salmonella*. The analysis has provided new hypotheses for future experiments to study this host defense mechanism.

Various groups have proposed theoretical concepts to analyze the knockout behavior of metabolic networks and gene regulatory networks see, for example, Steinway *et al.,* 2015, Padwal *et al.,* 2014. For logical models, software tools exist, e.g., GINsim (Naldi *et al.,* 2009), BooleanNet (Müssel *et al.,* 2010), CellNetAnalyzer (Klamt *et al.,* 2007). These tools do not consider Manatee invariants and / or the steady-state condition. To fill this gap, we implemented a stand-alone, easy-to-use tool for *in silico* knockout analyses, in particular for for experimentally working scientists. Here, we describe the implementation of a new method - isiKnock (**in si**lico **knock**out) - a software to systematically perform and visualize *in silico* knockouts in mathematical models of biochemical systems. Petri net (PN) formalism serves as modeling language since they were successfully used in systems biology, for an overview see, for example, (Koch *et al.*, 2011). We consider signaling pathways to investigate possible signaling flows from receptor activation to the cell response, see Scheidel *et al.,* 2016 and Amstein *et al.,* 2017.

## 2 Features

*In silico* knockouts investigate the influence of disabled reactions, e.g., an impaired protein synthesis, represented by transitions of the PN, on the signaling pathway. The prediction of knockouts requires the computation of Manatee invariants, which represent combinations of transition invariants and reveal all possible signal flows from the signal initiation to the cell response (Amstein *et al.,* 2017). A non-functional or knocked out transition affects all Manatee invariants that contain this transition. The unaffected Manatee invariants describe the remaining functional parts of the network. The program automates the prediction of *in silico* knockout behavior for single as well as multiple knockouts.

isiKnock provides a graphical user interface (GUI), which enables the loading of PN models in different file formats, adjustment of the settings, and the simulation of the knockouts, see Fig. 1. We implemented the support of the file formats SBML (Hucka et al, 2003) and, for PN additionally, PNT, which represents a special PN format used in INA (Starke *et al*., 1999) and other PN-based tools. PN editors like MonaLisa (Einloft *et al*., 2013) give the possibility to construct a model or convert a SBML model into a PN.

**Fig 1.**
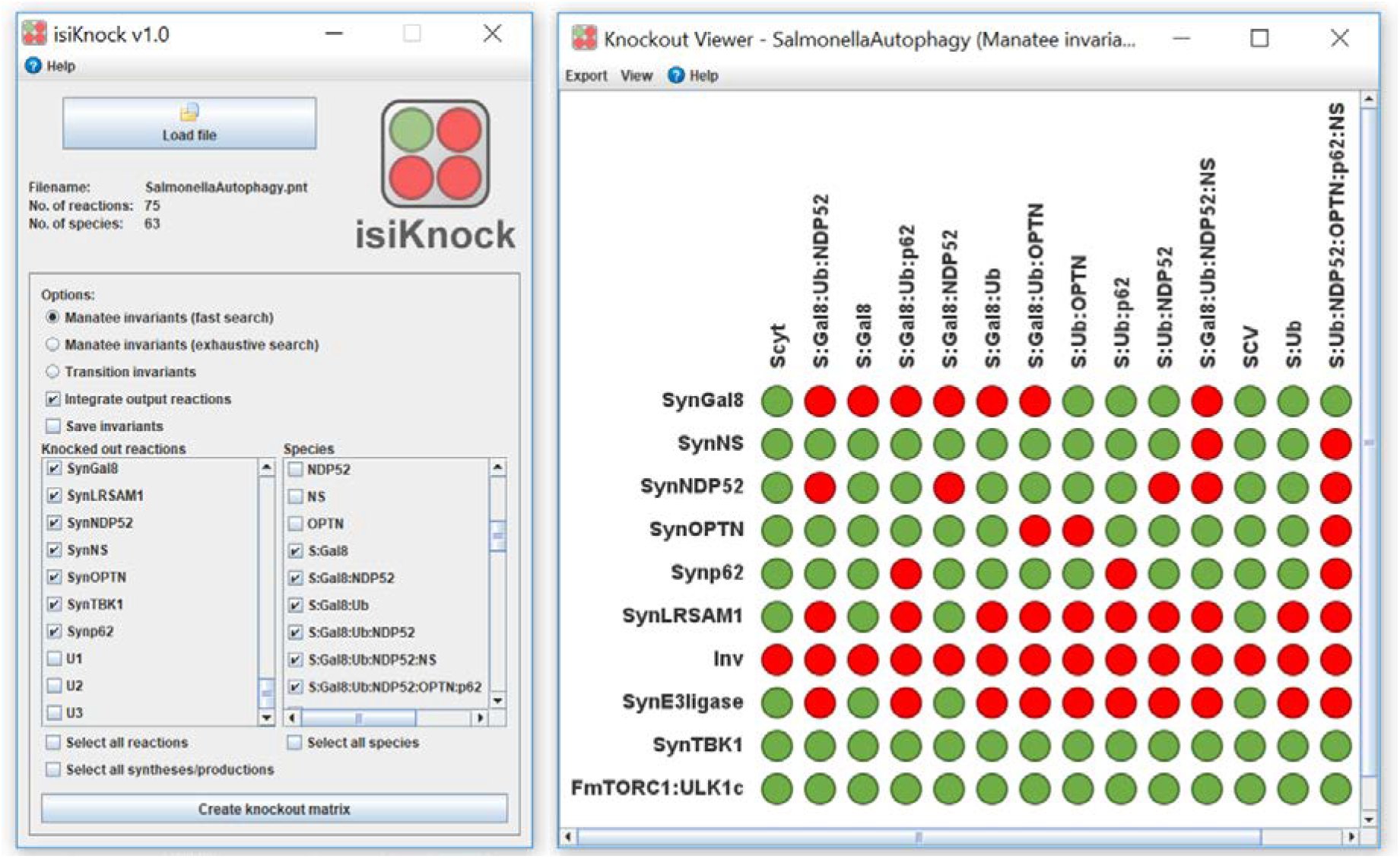
Graphical user interface of isiKnock. The left part shows the start panel. Here, the user can load the PN file and change settings. The button “Create knockout matrix” starts the computation of the knockout analysis. The right part depicts the knockout matrix for the network of the xenophagic capturing of *Salmonella* (Scheidel et al., 2016). The rows represent the knocked out reactions (transitions) and the columns the species (places). A green entry indicates no effect of the knockout, i.e., the formation of a protein complex is still functional at steady-state conditions. An entry is red if the species is affected by the knockout.

The GUI visualizes the predicted knockouts as color-coded matrix, red for affected and green for unaffected parts. The rows represent the knocked out reactions and the columns the species. The user can adjust the design of the matrix, e.g., the colors, the names of the reaction and species, or can change the order of the matrix entries (e.g., alphabetically). To detect groups of reactions that affect the system in the same way, e.g., syntheses of proteins with a similar function, or to reveal pathway components that are influenced by the system in the same way, e.g., groups of similarly regulated proteins, we can cluster matrix entries. There exists the possibility to export the matrix as PNG, SVG, or CSV file. A TXT file saves the transition invariants or Manatee invariants. For a detailed tutorial, see http://www.bioinformatik.uni-frankfurt.de/tools/isiKnock/.

In summary, we introduced a new open-source software for *in silico* knockouts, isiKnock. *In silico* knockouts can be easily conducted, the results are visualized and can be exported into several formats. The examination of the network behavior to perturbations of specific proteins enables to address important issues, such as dependencies of biological pathways, identification of potential targets for drug treatment and predictions of unknown effects of perturbations.

## Funding

This work was supported by the LOEWE program Ubiquitin Networks (Ub-Net) of the State of Hesse (Germany) [20120712/B4] and by the Cluster of Excellence “Macromolecular Complexes” of the Goethe-University of Frankfurt am Main [3212070002/TP2].

## Conflict of Interest

none declared.

